# Mutual information of high-dimensional random variables: estimation by frontier mutual information

**DOI:** 10.1101/2024.12.29.630641

**Authors:** Tsutomu Mori, Takashi Kawamura

## Abstract

Many natural sources of information, including genes, yield intricate datasets characterized by high-dimensional random variables. However, the computational constraints and information loss have often limited the accuracy of mutual information (MI) computations in such datasets. To address these limitations, we introduce a novel metric, micromutual information, which measures the information exchange at each cell level within high-dimensional contingency tables. This methodology represents an extension of our previous techniques and employs a linear index approach. The method simplifies complex, high-dimensional genetic data into a one-dimensional format, thereby improving computational efficiency while preserving the intricate structure of gene interactions. Theorems are developed which demonstrate how the sum of micromutual information asymptotically converges to the total MI for multidimensional variables. Our findings indicate that the maximum value of micromutual information, termed *MI*_*front*_, adheres to an extreme value distribution. The observation of *MI*_*front*_ provides a streamlined approach to estimating the total MI, due to the simplicity of measuring the micromutual information of just one cell. This approach has the potential to improve data analysis in genomics and other fields that deal with multidimensional information.

## I. Introduction

**T**He intricate dynamics of gene interactions present a significant challenge to contemporary research, both in the fields of biology and information theory. While traditional feature extraction methods such as kernel density estimation (KDE) [8] and conditional mutual information (CMI) [17] are useful in a variety of scientific disciplines, they are often inadequate for the high-dimensional nature of genetic interactions. These methods are challenged by computational complexity and information loss in high-dimensional settings. Furthermore, they fail to account for the specific and dynamic nature of gene interactions, which are influenced by temporal, spatial, and physiological conditions. This makes it difficult to capture their transient and context-specific behaviors.

To address these challenges, this study proposes a novel metric, micromutual information (*MI*_*kl*_). *MI*_*kl*_ is designed to measure information exchange at the level of individual cells within high-dimensional contingency tables, with the potential to capture subtle interactions between genes that may be missed by conventional methods. In addition, the use of a linear indexing method improves computational efficiency while preserving the complex structure of gene interactions, providing a basis for detailed analysis of specific interactions.

By integrating these methods, we developed the “*ab initio* Genetic Orbital (GO) Method” [9], which aims to predict gene functions. This method postulates that genetic information and intergenic interactions evolve through stochastic processes, starting from an initial state where mutual information (MI) is zero. This assumption justifies the use of a uniform distribution as the initial condition for biological information. Moreover, we derived the micromutual information summation theorem, which states that the sum of *MI*_*kl*_ asymptotically converges to the total MI of multidimensional random variables. By applying these principles, the GO method facilitates the estimation of MI between genes without the need for experiments. Our approach offers a promising avenue for analyzing specific and context-dependent interactions in genetic data, complementing conventional methods.

Our findings indicate that the largest value of micromutual information (*MI*_*front*_) is best described by an extreme value distribution. Consequently, observing *MI*_*front*_ offers a streamlined approach to estimating total MI, primarily due to the simplicity of measuring just one cell s *MI*_*kl*_. In contrast to conventional methodologies that aggregate mutual information across multiple dimensions, our approach emphasizes the examination of micro-level mutual information *MI*_*kl*_ and its maximal value *MI*_*front*_, which may otherwise be obscured by aggregating information across diverse dimensions. This targeted analysis enables the precise quantification of gene interactions under a specific set of conditions, thus identifying the most influential relationships that drive biological processes or pathological conditions.

As previously demonstrated in *Jxiv* 2023 [9], the GO method successfully applied the above rationale to the analysis of gene networks from a large-scale database. In that research, we identified *KYNU* as a critical gene involved in tumor immunity, thereby demonstrating the practicality and effectiveness of focusing on *MI*_*front*_. In this paper, we provide a comprehensive examination of the methodologies presented in the aforementioned paper, thereby emphasizing their significance in overcoming the conventional limitations associated with the analysis of high-dimensional genetic datasets.

Prior to addressing the treatment of high-dimensional random variables, it is necessary to consider the case of one-dimensional random variables that assume a multitude of discrete values. Let *X* and *Y* be discrete random variables, taking on values between *X*_1_ and *X*_*m*_ and *Y*_1_ and *Y*_*n*_, respectively. In this context, *m* and *n* are both natural numbers. We consider an *m* × *n* contingency table with respect to *X* and *Y* (see Fig. 1a). The joint probability of the cell (*k, l*) is represented by *p*(*X*_*k*_, *Y*_*l*_), whereas the marginal probabilities of the *k*-th column and the *l*-th row are given by *p*(*X*_*k*_) and *p*(*Y*_*l*_), respectively. The mutual information (MI) of *X* and *Y*, denoted *MI*, is then defined as follows [13].

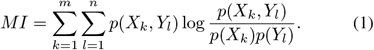

**Fig. 1.**
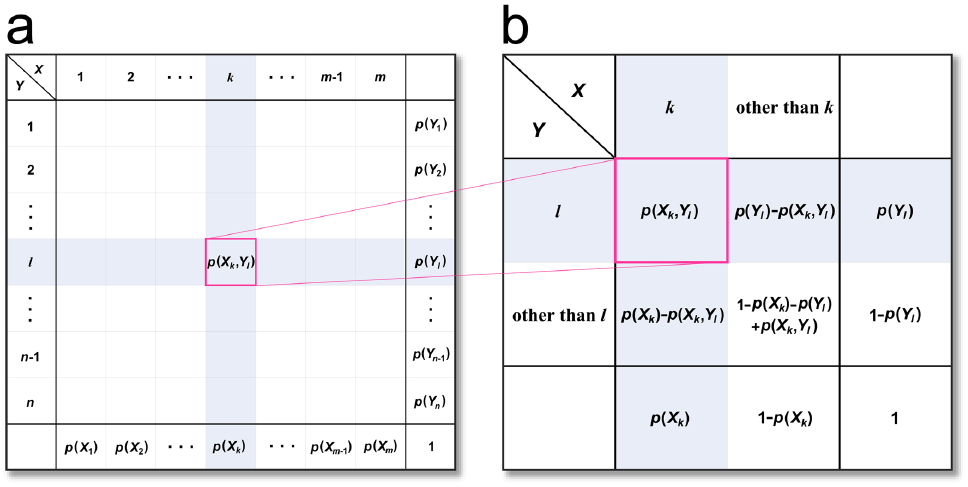
Micromutual information of a contingency table. (a) An m ×n contingency table with respect to the random variables X and Y. (b) A 2× 2 contingency table with respect to the cell (k, l) of (a).

The MI is defined as the mean information about variable *Y* given by variable *X*. It can also be expressed as the mean information about variable *X* given by variable *Y*. Furthermore, the MI is always non-negative. The MI has been identified as a prominent measure of the interdependence between *X* and *Y* in numerous studies, including those referenced in [1, 6, 11, 16]. Nevertheless, the calculation of the MI of multidimensional random variables has proven to be a challenging endeavor.

In order to address the limitations of applying *MI* to high-dimensional information sources, we introduce a new metric, *MI*_*kl*_, which is defined for a cell (*k, l*) of the original contingency table [9] as follows:

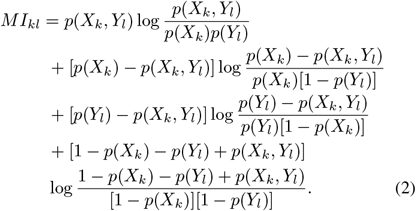

In this context, *MI*_*kl*_ represents the MI of the 2 × 2 contingency table that is derived from the original *m* × *n* table. This aforementioned table illustrates the occurrences of *X* when taking *X*_*k*_ and *Y* when taking *Y*_*l*_, as demonstrated in Figure 1(b). This transformation quantifies the specific information exchange between *X* and *Y* under these particular conditions, thereby simplifying the analysis of the dataset by focusing on key interactions and reorganizing the data to highlight relevant events. The reduced table allows for the capture of specific interactions under specific combinations of conditions, thereby facilitating the handling of complex multidimensional information in a more accessible format.

As a subsequent step, we analyze the interrelationship between the original MI and the sum of *MI*_*kl*_, where *k* and *l* range from 1 to *m* and from 1 to *n*, respectively. The following Theorem 1 asserts that as the values of *m* and *n* approach infinity, the sum of *MI*_*kl*_ converges to *MI*. In light of this, the present paper addresses the subject of asymptotic theory, which posits that when *m* and *n* approach infinity, the MI can be calculated as the total sum of *MI*_*kl*_ observed at each state. The conclusions of Theorem 1 are applicable to any one-dimensional random variables, regardless of the complexity of their distributions.

### Theorem 1

*Let X and Y be discrete random variables, which take X*_1_ *to X*_*m*_ *and Y*_1_ *to Y*_*n*_, *respectively. Let MI be the MI of X and Y*, *and let MI*_*kl*_ *be the micromutual information with respect to the cell* (*k, l*), *where k and l take 1 to m and 1 to n, respectively. Then the total sum of MI*_*kl*_ *converges to MI, as m and n tend to infinity*.

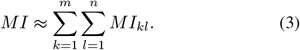

In this paper, we designate this formula as the micromutual information summation theorem for one-dimensional random variables. As *m* and *n* approach infinity, this theorem implies the emergence of additivity of *MI*_*kl*_, which also signifies that each *MI*_*kl*_ tends towards mutual independence.

The aforementioned arguments are also applicable to multi-dimensional random variables, which represent the primary focus of this paper. The MI of multidimensional random variables can be obtained by employing the linear index method, as outlined in reference [9]. This method entails transforming the variables into a one-dimensional format, calculating their *MI*_*kl*_, and subsequently taking the sum to ascertain the MI. In Section III, we prove Theorem 2, which is the micromutual information summation theorem that is applicable to multidimensional random variables.

In our preceding paper, we demonstrated that MI exhibits distinctive properties from a statistical and probabilistic point of view [10]. In this paper, we present an efficient method for estimating MI that exploits these properties. In the absence of prior information regarding the distribution of random variables, it is reasonable to assume that they follow a uniform distribution according to the maximum entropy principle [3]. As previously demonstrated [10], the probability distribution of MI for random variables *X* and *Y* following uniform distributions can be expressed as *P* = *e*^*−MI*^.

In light of this, we demonstrate in Theorem 4 that when *X* and *Y* follow a uniform distribution and *m* and *n* approach infinity, *MI*_*kl*_ also follows an exponential distribution. Furthermore, the maximum value of *MI*_*kl*_, denoted *MI*_*front*_, follows a general extreme value distribution. By employing these probability distributions in conjunction with the micromutual information summation theorems, it is possible to estimate *MI* from the measurement of *MI*_*front*_ of only one cell. Accordingly, this methodology allows the efficient estimation of the MI of multidimensional random variables.

Moreover, we examine another aspect of the relationship between MI and statistics. As has been demonstrated previously, MI is asymptotically equivalent to the *P*-value, which represents a principal statistical measure [10]. For a statistical contingency table of sample size *N*, the significant probability, or *P*-value, of the MI can be derived if *N* is sufficiently large [10]. The *P*-value is calculated using the following formula *P* = *e*^*−N·MI*^. In order to ascertain the MI, it is essential to utilize an *m* × *n* contingency table of joint probability distributions, as shown illustrated in Figure 1(a). This aforementioned table can be obtained by dividing the cell frequency by *N*, where *N* represents the sample size of the contingency table. As in statistics, this principle can be applied to perform meta-analyses that combine multiple independent datasets in order to accurately calculate the MI between random variables.

In comparison to the conventional methods for estimating the MI of one-dimensional random variables, our proposed GO method exhibits notable distinctions. Among the afore-mentioned traditional methods, the KDE method [8] and the *k*-th nearest neighbor method [7] are the most commonly employed. The former method estimates the probability density function as a superposition of Gaussian kernels. The latter method identifies the *k*-th nearest neighbors of each data point and creates local bins in the vicinity of each point. In contrast, our approach is markedly distinct from these existing methods. Furthermore, it is capable of accommodating multidimensional random variables and computing the *MI*_*kl*_ for a 2 × 2 contingency table derived from the original table with fixed bins. One of the advantages of our method over alternative approaches is its capacity to discern the combination of states in which random variables interact most significantly. In particular, our method is effective for biological datasets because a significant proportion of biological variables can be considered as multidimensional random variables.

In summary, the *ab initio* GO method not only refines the feature extraction and dimension reduction techniques for complex genomic datasets, but also provides a novel framework for understanding the intricate dependencies within gene networks. This approach represents a significant departure from traditional methods and provides a more profound understanding of the biological significance of gene interactions.

The following is a description of the organization of this paper. In Section II, the proof of Theorem 1 is presented. The proof of Theorem 2 for multidimensional random variables is presented in Section III. In Section IV, we investigate the impact of random variable dimensionality on MI and prove Theorem 3. Section V delineates the *P*-value associated with the estimation of the MI of multidimensional random variables and its correlation with the statistical *P*-value of a contingency table of sample size *N*. Section VI substantiates Theorem 4, which postulates that the micromutual information *MI*_*kl*_ exhibits an exponential distribution in its asymptotic behavior. Section VII is devoted to an examination of the properties of frontier mutual information, denoted *MI*_*front*_. Section VIII presents a series of numerical simulations. Section IX comprises the conclusion and discussion.

## II. Proof of Theorem 1

According to (1) and (2), the difference between *MI* and the sum of *MI*_*kl*_ over all the cells is given by

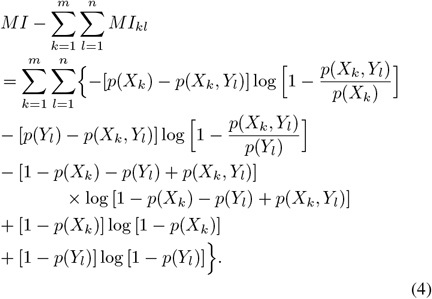

Next, we divide each cell of the original contingency table into *h* equal parts for both row and column directions. General divisions of the cells can be reduced to this division. If we take the sum of *MI*_*kl*_ over all the divided cells, then we get

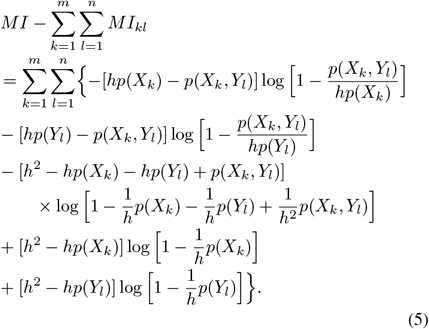

Using L’Hôpital’s theorem, we calculate the limit of each term on the right-hand side of (5) as *h* approaches infinity. The limits of the first and second terms are both *p*(*X*_*k*_, *Y*_*l*_). In fact, the limit of the first term of (5) is computed as

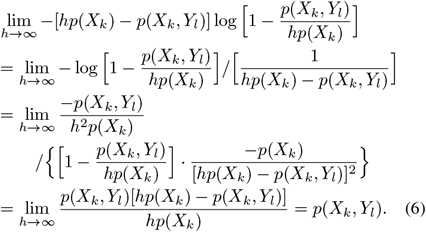

Meanwhile, the third term approaches

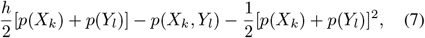

and the fourth and the fifth terms respectively approach

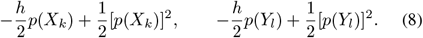

If we add up these limits, keeping in mind that the sum of *p*(*X*_*k*_, *Y*_*l*_), that of *p*(*X*_*k*_), and that of *p*(*Y*_*l*_) is equal to 1, we can see that the limit of the right-hand side of (5) is zero.

Actually,

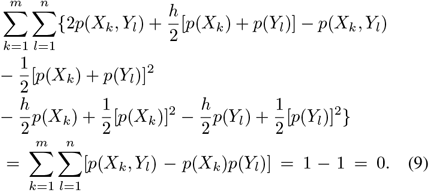

This shows that the right side of (5) is approaching zero.

Thus, when *m* and *n* tend to infinity,

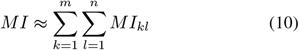

holds. This proves Theorem 1. □

## III. Application to multidimensional random variables

This section presents a method for applying Theorem 1 to the computation of the MI of multidimensional random variables. For illustration, let *Z* be a *d*-dimensional discrete random variable, where *Z* = (*W*_1_, *W*_2_, …, *W*_*d*_). Each of the *W*_*i*_(1≤ *i* ≤*d*) is a one-dimensional discrete random variable, with an index set ranging from 1 to *m*_*i*_ [9]. By introducing a linear index, designated as *s*, to order the configurations of *Z* in a lexicographic manner, we transform *Z* into a one-dimensional variable designated as *Z*_*s*_, which ranges from *Z*_1_ to *Z*_*t*_ with 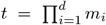. This reorganization permits the treatment of *Z* as either variable *X* or *Y* in the context of Theorem 1, thereby facilitating the derivation of Theorem 2.

### Theorem 2

*Let Z and U be multidimensional discrete random variables. We define Z*_*s*_ *and U*_*q*_ *as the one-dimensional random variables transformed from Z and U by the linear indices s and q, which run from 1 to t and from 1 to v, respectively. Let MI*_*sq*_ *be the micromutual information with respect to the cell* (*s, q*) *of the transformed contingency table. Then the overall sum of MI*_*sq*_ *approaches the MI of Z and U*, *MI, as t and v increase without bound*.

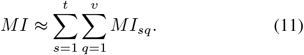

In this paper, we designate this formula as the micromutual information summation theorem for multidimensional random variables. By transforming multidimensional random variables into one-dimensional ones, it is feasible to calculate *MI*_*sq*_ and MI of multidimensional random variables, thereby substantiating the veracity of Theorem 2. As *t* and *v* approach infinity, Theorem 2 suggests the additivity of *MI*_*sq*_, which signifies that each *MI*_*sq*_ approaches mutual independence.

## IV. Effect of dimension on MI

Subsequently, the impact of the dimension *d* is determined. For two *d*-dimensional random variables, it is postulated that their components assume *m*_0_ and *n*_0_ values, respectively. The linear index is then employed to transform the variables into one-dimensional random variables, each of which assumes *m* = *m*_0_^*d*^ and *n* = *n*_0_^*d*^ discrete values, respectively. Then *MI* is expressed by using the mean ⟨*MI*_*sq*_⟩ of *MI*_*sq*_ as (*m*_0_*n*_0_)^*d*^ ⟨*MI*_*sq*_⟩, since the total number of the cells in the contingency table is (*m*_0_*n*_0_)^*d*^. This results in Theorem 3.

### Theorem 3

*Let Z and U be d-dimensional discrete random variables. We assume that their components take m*_0_ *and n*_0_ *values, respectively. The one-dimensional random variables Z*_*s*_ *and U*_*q*_ *are defined as transformations of Z and U*, *respectively, by the linear indices s and q. Let MI*_*sq*_ *be the micromutual information with respect to the cell* (*s, q*) *of the transformed contingency table. As the dimension of the random variables, d, tends to infinity, the MI is expressed as follows:*

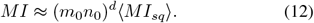

The augmentation of the number of cells in the contingency table signifies the increasing intricacy of the interactions between the random variables. To illustrate, if *m*_0_ = *n*_0_ = 5 and *d* = 4, then *MI* is approximately 3.9 × 10^5^ times greater than ⟨*MI*_*sq*_⟩. Similar considerations can also be applied to infinite-dimensional discrete random variables, that is, when *d* tends to infinity. In light of the plethora of ways in which information is conveyed by a gene, it can be regarded as an infinite-dimensional discrete random variable [9]. As *d* tends to infinity, the maximum of *MI* for genes increases rapidly as a function of *m*_0_, *n*_0_, and *d*. This finding indicates that the interactions between genes become more diverse, thereby facilitating a more precise exchange of information between them. It seems reasonable to posit that these effects have provided biological organisms with evolutionary advantages.

## V. Probability-statistical implications of MI of high-dimensional random variables

This section examines the probability-statistical implications of the MI of very high-dimensional random variables, as analyzed by contingency tables. The law of large numbers guarantees that as the sample size *N* grows, the relative frequency observed in each cell of a contingency table approaches the true joint probability. This convergence permits the approximation of the true MI by substituting the aforementioned relative frequencies into the definition of *MI* (1) when *N* is large. Consequently, a statistical contingency table becomes asymptotically equivalent to a contingency table based on the joint probability (Fig. 1a). When this equivalence is satisfied, the methods of probability statistics may be applied to *MI*, as will be described in the following text.

The first case examines the joint probabilities in the absence of *N*. The objective is to investigate the probability distribution of *MI* in the context of a lack of prior information regarding the distributions of the random variables. Given the inherent randomness of genetic information, it is reasonable to assume a uniform distribution in accordance with the maximum entropy principle. When *X* and *Y* are distributed uniformly according to this supposition, their MI is distributed exponentially as follows, as demonstrated in [10].

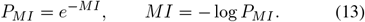

The left formula also represents the significance probability (*P*-value), thereby underscoring the augmented probabilistic interdependence as the magnitude of *MI* increases. As the dimension *d* of the contingency table or the category counts (*m* and *n*) increase, the maximum value of *MI* also increases. This is equivalent to a decrease in the minimum *P*-value, allowing a possible strong dependency as the size of the contingency table grows.

To illustrate, in the case presented in Section IV where *m*_0_ = *n*_0_ = 5 and *d* = 4, if ⟨*MI*_*sq*_⟩ = 10^*−*5^, then *MI* ≈ 3.9, and *P*_*MI*_ ≈ 0.020, as derived from (13), indicating a significant interdependence. In this case, if *m*_0_, *n*_0_, and *d* are each increased by a factor of two, the maximum *MI* increases by a factor of 1.10 ×10^12^, while the minimum *P*-value decreases by a factor of 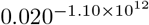. Thus, as *m*_0_, *n*_0_, and particularly *d* increase, the maximum MI increases considerably, while the minimum *P*-value decreases dramatically. This result underscores the necessity for a combination of statistical and information-theoretic techniques to characterize the interactions between high-dimensional random variables, such as genetic data. Moreover, it illustrates the importance of (13) in meeting these requirements.

The next case study examines the frequencies and *N*. To investigate this, we must consider a statistical contingency table of a sufficiently large sample size *N* about *X* and *Y*, which follows a uniform distribution. In this context, the probability *P*_*N·MI*_ that the MI of *X* and *Y* becomes *MI* is represented as follows, as proved in [10].

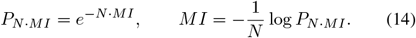

In this context, *P*_*N·MI*_ also represents the *P*-value that the MI is equal to or greater than *MI*. When these formulas are applied to the above example, if ⟨*MI*_*sq*_⟩ = 10^*−*5^*/N*, then *MI* ≈ 3.9*/N*, and *P*_*N·MI*_ ≈ 0.020 from (14), indicating a significant interdependence. Accordingly, the statistical *P*-value of a high-dimensional contingency table can be readily calculated by summing *MI*_*sq*_ from Theorem 2. Therefore, as *d, m* and *n* increase, the maximum MI increases, the minimum *P*-value decreases, and the interdependence between the random variables may potentially become significant.

These results demonstrate how statistical and information-theoretic methods converge to provide significant insights. Increasing the dimensions and categories of contingency tables not only increases the maximum MI but also gives knowledge of complex dependencies in genetic interactions. Subsequent sections will utilize the above findings to derive the probability distribution of *MI*_*kl*_ and develop methods for meta-analysis.

## VI. Exponential distribution of micromutual information

This section presents a proof of Theorem 4, which demonstrates that *MI*_*kl*_ follows an exponential distribution, when *m, n*, and *N* are sufficiently large. To illustrate, consider extensive databases such as The Cancer Genome Atlas (TCGA) [15], which is presumed to satisfy the conditions of this theorem. TCGA is a comprehensive database project that collects and analyzes genomic data from various types of cancer. The database provides researchers with invaluable resources for understanding the molecular basis of cancer, offering data such as genomic sequences, gene expression, copy number variation, and epigenetic information. Given its substantial sample size (*m, n*, and *N*), TCGA represents an appropriate illustration of Theorem 4.

Prior to presenting Theorem 4, it is necessary to elucidate the significance of large *m, n*, and *N*. In the context of complex information sources, such as genetic datasets where each dimension can encompass a vast number of categories, it is of paramount importance to guarantee that the true value of *MI*_*kl*_ is accurately determined. In order to address the inherent complexity of such data, it is necessary to verify that the categories *m* and *n* of the contingency table are sufficiently large. Concurrently, a large sample size *N* is essential to confirm that each cell in the table is adequately populated. The stability of the estimates of *MI*_*kl*_ across the table ensures that *MI*_*kl*_ is independent of the other cells and corresponds to a true probability distribution. Consequently, *MI*_*kl*_ accurately reflects the level of information exchange between the variables. A large *N* contributes to the reliability of the statistical inference and facilitates the reliable interpretation of the interactions within the high-dimensional genetic information space. These considerations lead to the statement of Theorem 4.

### Theorem 4

*Let X and Y be discrete random variables, taking on values between X*_1_ *and X*_*m*_ *and Y*_1_ *and Y*_*n*_, *respectively, with the observed values determined by a random process. Let MI represent the mutual information between variables X and Y*. *Additionally, let MI*_*kl*_ *denote the micromutual information with respect to the cell* (*k, l*), *where k and l take on values between 1 and m and 1 and n, respectively. As the values of m and n approach infinity, the MI*_*kl*_ *variable follows an exponential distribution, represented by the probability density function p*(*x*) *as*

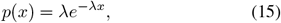

*where x* = *MI*_*kl*_ *and λ* = ⟨*MI*_*kl*_⟩^*−*1^.

*Proof of Theorem 4*. The proof of Theorem 4 is presented from two aspects. The initial aspect employs the tenets of Theorem

1. For sufficiently large values of *m* and *n*,

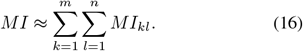

It can be seen that each summand on the right-hand side of equation (1), which defines *MI*, is identical to the first term on the right-hand side of equation (2), which defines *MI*_*kl*_. The sum of *MI*_*kl*_ with respect to *k* and *l* is calculated in accordance with the methodology outlined in (16). Consequently, the aggregate of the second, third, and fourth terms on the right-hand side of (2) with respect to *k* and *l* converges to zero. In other words,

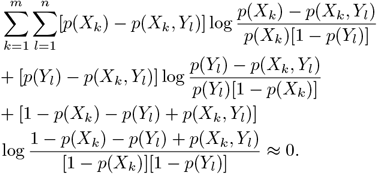

This indicates that as *m* and *n* approach infinity, the initial term on the right-hand side of (2) becomes less contingent on the joint probabilities of the other cells. Consequently, in a large contingency table, each *MI*_*kl*_ becomes increasingly independent of the micromutual information with respect to the other cells on average.

In contrast, the second aspect employs the principle of maximum entropy. As the values of *m* and *n* increase, the distributions of *X* and *Y* become more complex. Therefore, our understanding of these variables is limited, and the assumption of a uniform distribution is the optimal choice in accordance with the principle of maximum entropy. In this case, the values of *X* and *Y* are assumed to be drawn at random from the distributions defined by *X*_*k*_ and *Y*_*l*_, respectively. Meanwhile, as *N* tends to infinity, the relative frequencies of the *k*-th column, the *l*-th row and the cells respectively approach 1*/m*, 1*/n* and 1*/*(*mn*) according to the law of large numbers. Consequently, these values become less dependent on the relative frequencies of the other columns, rows and cells. Therefore, as *m, n* and *N* approach infinity, the *MI*_*kl*_ with respect to a given cell is largely independent of the mutual information with respect to the other cells.

Based on the two aspects mentioned above, we have arrived at the notion that each *MI*_*kl*_ is asymptotically independent. Consequently, from a microscopic point of view, when *m, n* and *N* are sufficiently large, we can apply (13) to individual *MI*_*kl*_, and *MI*_*kl*_ thus follows an exponential distribution. From a macroscopic perspective, the MI is constituted by ⟨*MI*_*kl*_⟩, and the mean value of *MI*_*kl*_ is calculated as

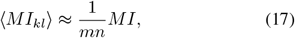

in accordance with the law of total expectation. Given that *MI* is presumed to adhere to an exponential distribution, it is similarly expected that *MI*_*kl*_ will also follow an exponential distribution. Accordingly, the dual perspective, microscopic and macroscopic, leads to the same conclusion.

In light of the above-mentioned circumstances, the probability density function of *MI*_*kl*_ is represented as follows:

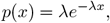

where *λ* = ⟨*MI*_*kl*_⟩ ^*−*1^. In this case, *p*(*x*) is normalized in such a way that

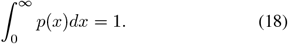

Thus Theorem 4 is proved. □

## VII. Frontier mutual information

### A. Definition and properties of frontier mutual information

Theorem 4 shows that micromutual information follows the exponential distribution (15). Therefore, the largest *MI*_*kl*_ is critical in that *X* and *Y* interact and transmit information to each other most strongly under this condition. In this section, we define the frontier mutual information, *MI*_*front*_, as the largest micromutual information and consider its properties.

*MI*_*front*_ has variations that consist of two parts. The first part is due to the sampling from the exponential distribution expected in the infinite contingency table. The second part is the shift from the exponential distribution due to the variations of *p*(*X*_*k*_, *Y*_*l*_), *p*(*X*_*k*_) and *p*(*Y*_*l*_). Since *MI*_*front*_ is the largest of all *MI*_*kl*_ of the finite contingency table, it is less independent of the micromutual information with respect to the other cells. It then tends to deviate from the exponential distribution.

First, we estimate *MI* from *MI*_*front*_, assuming that *MI*_*kl*_ follows the exponential distribution. From (15), if we let *j* be the rank of the magnitude of *MI*_*kl*_, say; *x*, then; for sufficiently large *m, n*, and *N*,

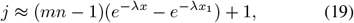

where *x*_1_ is the maximum *MI*_*kl*_.

Hence, *x* is approximately represented as a logarithmic function of *j* as

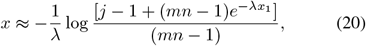

and

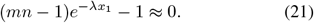

It follows that *MI*_*front*_ = max(*MI*_*kl*_) is related to *MI* as

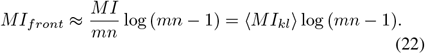

Thus, *MI*_*front*_ is approximately proportional to both the total *MI* and the mean of *MI*_*kl*_. Conversely, if *MI*_*front*_ follows the exponential distribution and contains no errors from it, then the total *MI* is estimated from *MI*_*front*_ as

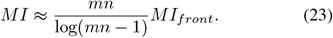

The variations of *MI*_*front*_ are discussed in the next subsection.

### B. Variations of frontier mutual information

In this subsection, we evaluate the two parts of the variations of *MI*_*front*_ described in the preceding subsection. Both of the two parts follow the distributions of maximum values. However, the types of distributions differ as follows.

First, the first part of the variation is estimated, which pertains to the maximum of the *MI*_*kl*_ samples. In accordance with the theory of order statistics [2], the distribution function of *MI*_*front*_ is given by

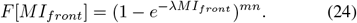

The mean of *MI*_*front*_, ⟨*MI*_*front*_⟩, is approximated as (*MI/mn*) log(*mn* − 1) by (23), which can also be derived from (24). The standa<u>r</u>d deviation *σ*[*MI*_*front*_] is approximately equal to 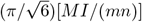. Accordingly, the 95% confidence interval for *MI*_*front*_ is given by

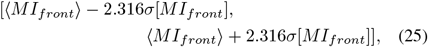

which is by substitution

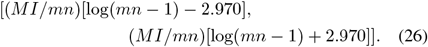

It is thus possible to estimate the 95% confidence interval of *MI* from *MI*_*front*_ as

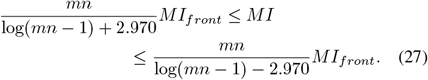

This interval is derived from the first part of the variation.

Next, an estimation is provided for the second part of the variation, which also affects *MI*_*front*_. The second part of the variation, Δ*MI*_*kl*_, of *MI*_*kl*_ can be derived from (2). As *N* approaches infinity, *p*(*X*_*k*_) and *p*(*Y*_*l*_) converge to 1*/m* and 1*/n*, respectively, resulting in the following:

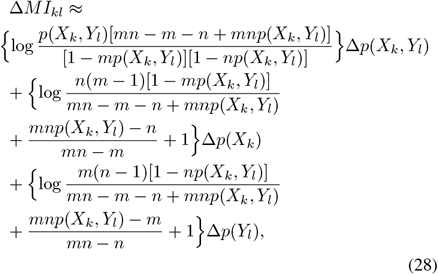

which is not zero in general. This demonstrates that *MI*_*kl*_ is subject to alteration in accordance with the fluctuations in joint probability, Δ*p*(*X*_*k*_, *Y*_*l*_), and the marginal probabilities, Δ*p*(*X*_*k*_) and Δ*p*(*Y*_*l*_). This constitutes the second part of the variation, whereby the distribution of *MI*_*front*_ differs from the exact exponential distribution.

Without the second part of the variation, the application of *MI*_*front*_ would enable the estimation of *MI* by means of the equation (23). However, the second part results in the observed *MI*_*front*_, which we designate as *MI*_*front*_*obs*_, being typically greater than the *MI*_*front*_ estimated by (22). We designate the latter as *MI*_*front*_*exp*_, where the suffix “ _*front exp*_” derives from the exponential distribution. To facilitate comparison, the ratio *MI*_*front*_*obs*_*/MI*_*front*_*exp*_ is evaluated. We subsequently demonstrate that it follows the generalized extreme value distribution [4]. This will be illustrated through the presentation of numerical simulations in the following section. The probability density function *f* (*y* |*µ, σ, ξ*) of the generalized extreme value distribution is represented as follows:

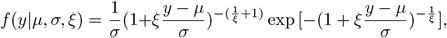

where *y* denotes *MI*_*front*_*obs*_*/MI*_*front*_*exp*_, *µ* is the location parameter, *σ* is the scale parameter, and *ξ* is the shape parameter. The results of the simulations indicate that the second part of the variation is greater than the first.

The confidence interval for *MI* is reevaluated by considering the variation of *MI*_*front_obs*_*/MI*_*front*_*exp*_. Let *MI* be the true value of MI, and let *MI*_*est*_ be the estimated MI from *MI*_*front_obs*_ by using (23) and dividing by *µ*. Under the assumption that *MI*_*front*_ follows an exponential distribution, as stated in (22), we obtain

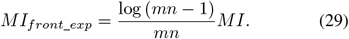

By definition, we get

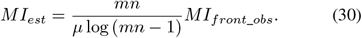

Therefore, we derive

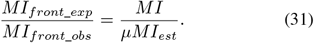

From (30) and (31), if the 95% confidence interval of *MI*_*front*_*exp*_*/MI*_*front*_*obs*_ is represented as

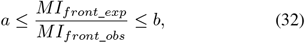

then the 95% confidence interval of *MI* is expressed as follows:

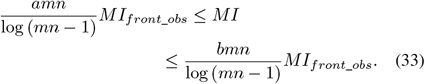

This formula (33) demonstrates that the measurement of *MI*_*front*_*obs*_ allows for the prediction of MI with a 95% confidence interval. In explicit terms, the values of *a* and *b* are represented as follows:

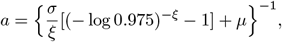

and

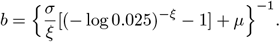

Therefore, despite the inherent difficulty in estimating *MI* between complex sources, this process is facilitated by focusing on *MI*_*front*_.

### C. Meta-analysis

The above results can be further applied to meta-analysis, which integrates data from multiple studies [12]. The objective of employing meta-analysis is to enhance the precision of estimating *MI* based on a single *MI*_*front*_*obs*_. In contrast to the conventional meta-analysis approach [12], which calculates the weighted average of *MI* without evaluating the *P*-value, our method considers the *P*-value of *MI* [10].

Let us consider the integration of *H* independent contingency tables with identical random variables and the same measurement errors, with sample size and MI given by *N*_*h*_ and *MI*_*h*_, respectively (where 1 ≤*h* ≤ *H*). In consideration of the independence of the tables, the composite *P*-value can be represented using (14) as

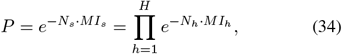

where the total sample size 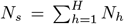 is sufficiently large, and *MI*_*s*_ is the true MI. Upon taking the logarithm of (34), *MI*_*s*_ is represented as

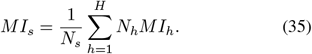

This suggests that the meta-analysis of MI can be conducted as *MI*_*s*_, the weighted average of *MI*_*h*_.

From (35), the standard error of *MI*_*s*_, Δ*MI*_*s*_, is expressed as follows:

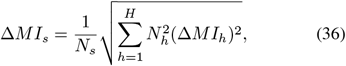

where Δ*MI*_*h*_ is the standard error of *MI*_*h*_. Moreover, the estimation of *MI*_*h*_ is derived from each *MI*_*front*_*obs*_ in accordance with the methodology delineated in the previous subsection. The error term Δ*MI*_*h*_ comprises two components: firstly, the error resulting from meta-analysis across the different tables; and secondly, the error resulting from the generalized extreme value distribution of *MI*_*front*_*obs*_*/MI*_*front*_*exp*_.

In particular, if *N*_*h*_ and Δ*MI*_*h*_ are constant across the different tables in (36), then

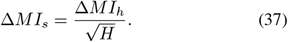

In this context, the value of Δ*MI*_*h*_, which encompasses both types of errors, remains constant. Consequently, the estimation of multiple *MI*_*h*_ improves the reliability of *MI*_*s*_ by augmenting the number of tables.

## VIII. Numerical simulations

To verify the validity of Theorems, a series of numerical simulations were conducted.

### A. Examples of Theorem 1

In this subsection, we present a series of numerical examples that illustrate the validity of Theorem 1. To begin, we present an illustrative example wherein *N* = 10^6^ points are randomly distributed in a contingency table of *m* = *n* = 10. Figure 2(a) depicts the resulting contingency table, while Figure 2(b) illustrates *MI*_*kl*_ with respect to the cells of the table. The maximum value of *MI*_*kl*_ was observed when *k* = 1 and *l* = 3, which corresponds to *MI*_*front*_, which characterizes the interactions between the random variables. The value of the cell in question did not exceed the mean value of 10,000. *MI* of the table shown in Figure 2(a) was 4.779×10^*−*5^, while the sum of *MI*_*kl*_ values across all 100 possible combinations of *k* and *l* in Figure 2(b) was 5.901× 10^*−*5^, which represents a 1.235-fold increase in MI. Therefore, when the parameters were set to *m* = *n* = 10, the summation of *MI*_*kl*_ for all possible combinations of *k* and *l* (i.e., 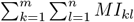) was found to be larger than *MI*, but still within a narrow range of its magnitude.

**Fig. 2.**
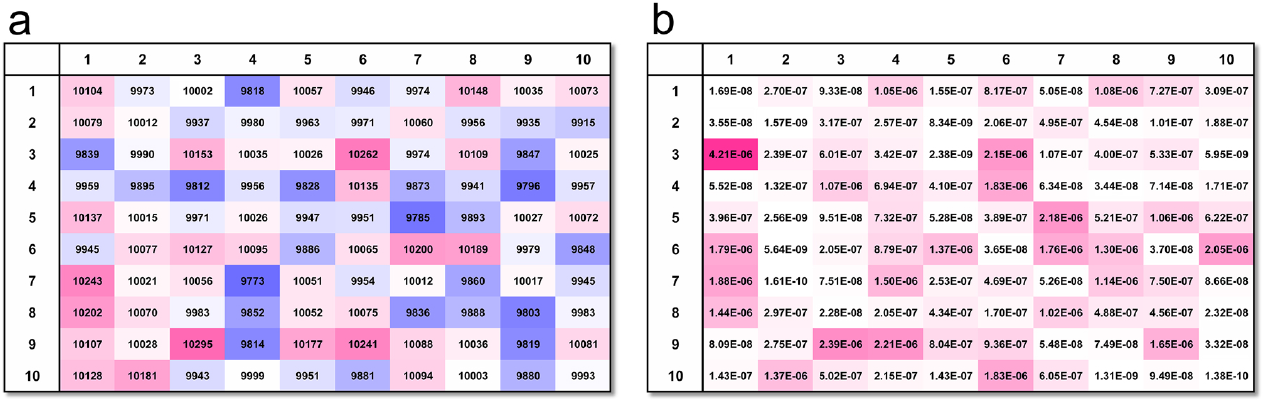
Numerical example of micromutual information. (a) Contingency table of m = n = 10 obtained by randomly distributing N = 10^6^ points. The red and blue cells have large and small numbers, respectively, according to the graduation. (b) MI_kl_ with respect to each cell (k, l) of (a). The unit of MI_kl_ is nat. The red cells have large MI_kl_ according to the graduation.

Next, we illustrate the reduction in the discrepancy between *MI* and the sum of *MI*_*kl*_ in accordance with the augmentation in *m* and *n*. We constructed contingency tables of *m* = *n* = 10 to 10^100^ by allocating one point per cell on average randomly and calculated *MI* and *MI*_*kl*_ (Fig. 3). The curves presented in Figure 3 which were generated with *m* = *n* = 10 to 10^4^ exhibit a convex upward trend. As *m* and *n* increase from 10 to 10^100^, the maximum of *MI* increases, 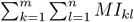 approaches *MI*, and the curve approaches a straight line when *m* = *n* = 10^100^, which supports Theorem 1.

**Fig. 3.**
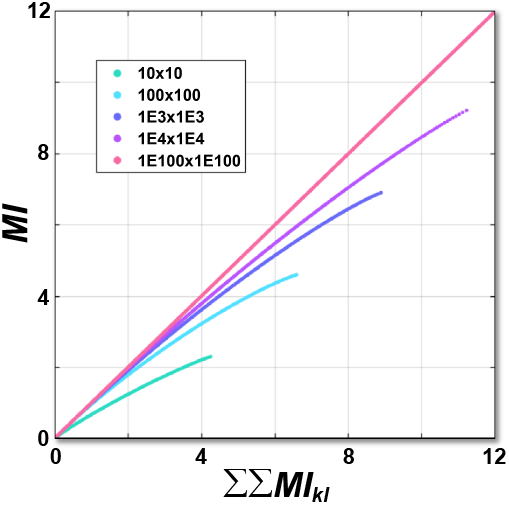
Relationship between MI and ∑ ∑ MI_kl_. MI and 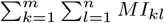 of contingency tables obtained randomly with m = n = 10 (green), 100 (light blue), 10^3^ (blue), 10^4^ (purple) or I0^100^ (red). The unit of MI and ∑ ∑ MI_kl_ is nat.

Moreover, the ratio max 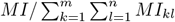 of the contingency tables was also calculated. In this context, max *MI* represents the maximum value of *MI* for each *m* = *n* (Fig. 4). As the values of *m* and *n* increase from 10 to 10^10^, the ratio max 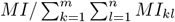 approaches one, thereby providing support for Theorem 1.

**Fig. 4.**
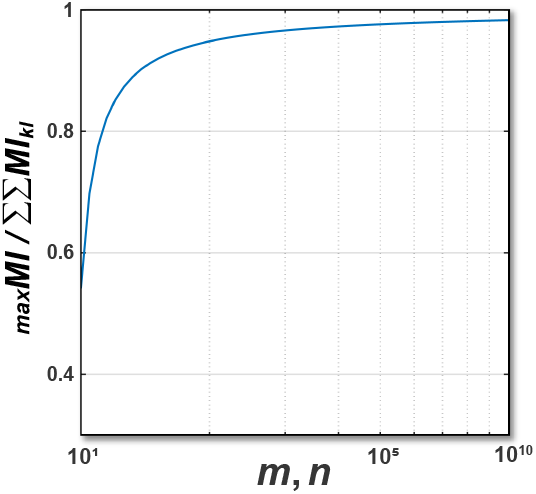
Ratio of MI to ∑ ∑ MI_kl_ for maximum MI contingency tables. 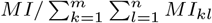 of the maximum contingency tables are plotted as a function of m = n. The abscissa is the common logarithm of m = n.

### B. Examples of Theorems 2 and 3

In order to prove Theorems 2 and 3, we have constructed the diagonal contingency tables of the multidimensional random variables *Z* and *U*, which take max *MI* for each *m*_0_ = *n*_0_ = 2 to 5 and *d* = 1 to 8. As *m*_0_, *n*_0_ and *d* increase, max *MI* (Fig. 5a) increases, particularly in a linear fashion with respect to *d*. Furthermore, max 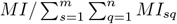 (Fig. 5b) also increases and approaches one, which lends support to Theorem 2. Moreover, max *MI/*⟨*MI*_*sq*_⟩ (Fig. 5c) also increases, approaches the number of cells (*m*_0_*n*_0_)^*d*^, and increases exponentially with respect to *d*, thereby confirming Theorem 3. The collective evidence presented in Figure 5(a), (b) and (c) suggests that an increase in dimension *d* induces an increase in the complexity of the interactions between the random variables.

**Fig. 5.**
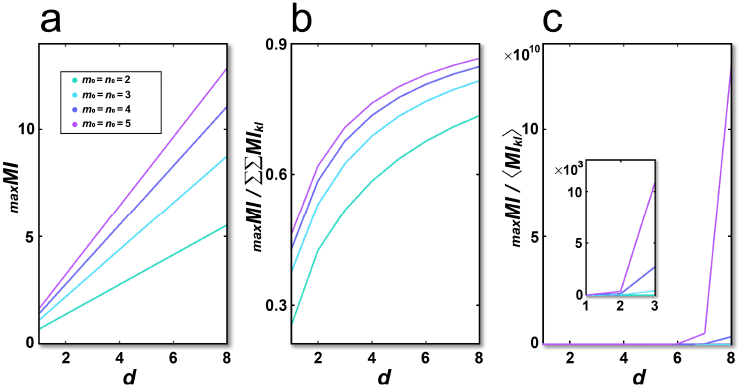
Maximum of MI as a function of the dimension d of random variables, (a) The maximum of MI (max MI) of the contingency tables, (b) max 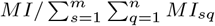. (c) max MI/ ⟨MI_sq_⟩. The inset depicts a magnification in the vicinity of the origin. The values of mo and no are 2 (blue), 3 (red), 4 (yellow) or 5 (purple), and d varies from 1 to 8. The unit of MI is nat.

### C. Examples of Theorem 4

To verify Theorem 4, we created contingency tables from *m* = *n* = 10 to 10000 by randomly distributing 10000 points per cell on average, and calculated *MI*_*kl*_ with respect to each cell of the tables. Figure 6(a), (b), (c) and (d) show the relative frequencies and their common logarithms of *MI*_*kl*_ of the tables with *m* = *n* =10, 100, 1000 and 10000, respectively. *MI*_*front*_ is indicated by the red arrows. These visualizations demonstrate that *MI*_*kl*_ almost follows the exponential distribution, and as *m* and *n* become larger, the distribution becomes more like the exact exponential distribution. This confirms the validity of Theorem 4 with numerical evidence.

**Fig. 6.**
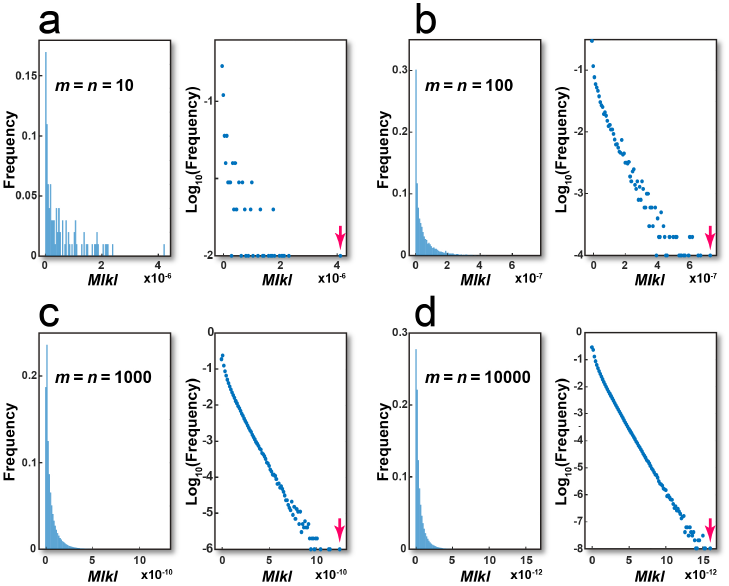
Exponential distribution of MI_kl_. (a) The relative frequency (left) and its common logarithm (right) of MI_kl_ of a contingency table with m = n = 10 obtained by distributing N = 10^6^ points randomly. (b) Those of a contingency table with m = n = 100 obtained by distributing N = 10^8^ points. (c) Those of a contingency table with m = n = 1000 obtained by distributing N = 10^10^ points. (d) Those of a contingency table with m = n = 10000 obtained by distributing N = 10^12^ points. MI_front_ is shown by the red arrows. The unit of MI_kl_ is nat.

In addition, as *m, n*, and *N* increase, the distribution of the relative frequencies of the cells approaches the normal distribution with mean 1*/*(*mn*) and variance 1*/*(*Nmn*) by the central limit theorem (Fig. 7). Similarly, those of the columns and rows approach the normal distributions with mean 1*/m* and 1*/n* and variance 1*/*(*Nm*) and 1*/*(*Nn*), respectively. The normal distribution has the maximum entropy of the distributions with known mean and variance. Therefore, these simulations realize the maximum entropy distributions, thus justifying the application of the maximum entropy principle.

**Fig. 7.**
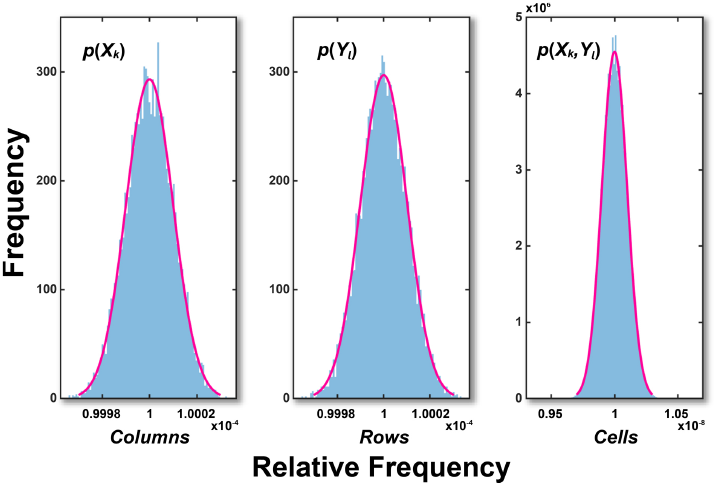
Histograms of relative frequencies. Distribution of the relative frequencies of the columns (left), rows (center) and cells (right) of the contingency table with m = n = 10000 obtained by putting N = 10^12^ points randomly. Red curves represent the fitting curves by the normal distributions. The mean and standard deviation are 1.000 × 10^−4^ and 1.005 × 10^−8^ (left), 1.000 × 10^−4^ and 1.000 × 10^−8^ (center) and 1.000 × 10^−8^ and 9.999 × 10^−11^ (right), respectively.

In Figure 6(a)-(d), the observed *MI*_*front*_*obs*_ was larger than the estimated *MI*_*front_exp*_ from (22). In order to consider their relationships and to deal with the actual larger MI, we performed simulations by making the contingency tables with *m* = *n* = 10 to 10000 one million times, the cells of which followed the normal distribution with mean 10000 and standard deviation 1000. Then, we confirmed that the values of *MI*_*front*_*obs*_*/MI*_*front*_*exp*_ follow the generalized extreme value distributions as mentioned above (Fig. 8). Since the shape parameters of the distributions are positive, they are Fréchet extreme value distributions. The 95% and 68% confidence intervals of *MI/MI*_*est*_ = *µMI*_*front*_*obs*_*/MI*_*front_exp*_ are shown in Figure 9, which shows that they become narrower as *m* increases.

**Fig. 8.**
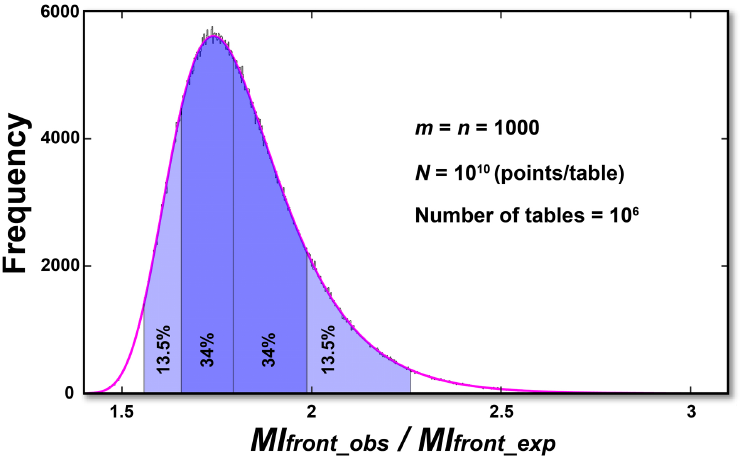
Generalized extreme value distribution of MI_front_obs_ / MI_front_exp_. Distribution of MI_front_obs_ / MI_front_exp_ from 10^6^ contingency tables with m = n = 1000, whose cells follow the normal distribution of mean 10000 and standard deviation 1000. The red curve is the fitted curve by the generalized extreme value distribution of the parameters µ = 1.742, σ = 0.1407 and ξ = 0.002590. The standard errors of µ, σand ξ are 1.57 × 10^−4^, 1.14 × 10^−4^ and 6.93 × 10^−4^, respectively.

**Fig. 9.**
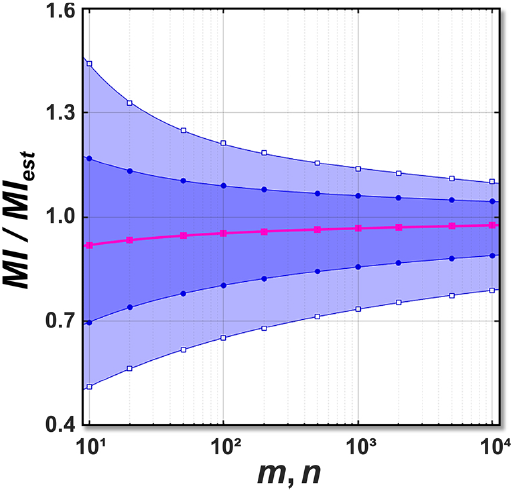
Lower and upper bounds of 95% and 68% confidence intervals of MI/MI_est_. The markers represent the lower and upper bounds of the 95% and 68% confidence intervals and the 50% cumulative probability of MI/MI_est_ calculated from 10^6^ contingency tables with m = n = 10 to 10000. The cells of the tables follow the normal distribution with a mean of 10000 and a standard deviation of 1000. The abscissa expresses m and n on a logarithmic scale. The light and dark blue areas represent the 95% and 68% confidence intervals, respectively. The red curve represents the 50% cumulative probability. From (31), µMI_front_exp_/MI_front_obs_ = MI/MI_est_.

Here, we describe numerical simulations to compare the 95% confidence intervals of *MI* given by (27) and (33). For example, we show a contingency table with *m* = *n* = 1000, whose cells follow a normal distribution with mean 10000 and standard deviation 1000. Its observed *MI* = 5.0 × 10^*−*3^ and *MI*_*front*_*obs*_ = 1.435 × 10^*−*7^. The 95% confidence interval from (27) using *MI*_*front*_*obs*_ corresponding to the sampling variation from the exponential distribution is calculated as

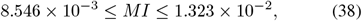

which does not contain the true *MI*. Therefore, it is not sufficient to consider only the exponential distribution.

Meanwhile, the 95% confidence interval of *MI* from (33) with calculated *a* = 0.3710 and *b* = 0.5751 is presented as

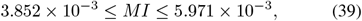

which contains the observed *MI*. Thus, the 95% confidence interval of the *MI* from (33) is more accurate than that from (27). Therefore, the inequalities (33) based on the generalized extreme value distribution are effective for calculating the confidence interval of the true *MI* from *MI*_*front*_*obs*_. Thus, the method of using *MI*_*front*_ is useful for computing *MI* of very high-dimensional random variables.

### D. Examples of meta-analysis

To demonstrate the effectiveness of our methodology in meta-analysis, we present a series of numerical examples (Fig. 10). The original contingency tables were created with *m* = *n* = 10, 100, 1000, or 10000, with cells following a normal distribution with a mean of 100000 and a standard deviation of 10000. These are designated as the mother contingency tables. By randomly sampling from the above mother contingency tables, the child contingency tables are constructed in such a way that their *m* and *n* are identical to those of the mother contingency tables, while their sample sizes are reduced by a factor of ten. The child contingency tables were replicated 10, 100 and 1000 times, respectively. Subsequently, the MI of the child contingency tables was then estimated from their *MI*_*front*_*obs*_ by using the formula provided in (30). This estimated MI is denoted as *MI*_*est_c*_. Then,

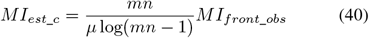

holds. Here, *µ* is the location parameter of the generalized extreme value distribution in Subsection VII.B. We attempted to estimate the MI of the mother contingency table, *MI*_*mother*_, from *MI*_*est_c*_ and calculated its ratio *MI*_*mother*_*/MI*_*est*_*c*_, which is denoted by *R*_*M*_.

**Fig. 10.**
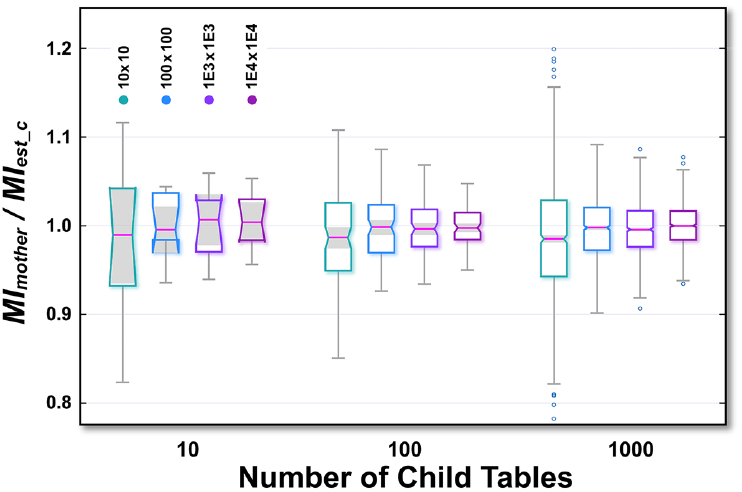
Box-whisker plots of R_M_ = MI_mother_/MI_est_c_ in meta-analysis. The mother contingency tables have m = n = 10 to 10000, whose cells of follow a normal distribution with mean 100000 and standard deviation 10000. The abscissa is the number of child tables. For each number of child tables, the first, second, third and fourth bars from the left represent R_M_ when m = 10, 100, 1000 and 10000, respectively. The shaded notches represent the 95% confidence intervals of the medians. The length of the shaded notch is equal to 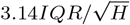, where IQR is the interquartile range and H is the number of child tables.

Figure 10 illustrates the box-whisker plots of *R*_*M*_. As *m* increases from 10 to 10000, and the number of child contingency tables increases from 10 to 1000, the median of *R*_*M*_ approaches 1. Additionally, the 95% confidence interval, indicated by the shaded notch, becomes narrower. This indicates that the standard error of the mean of *R*_*M*_ decreases, and the median of *R*_*M*_ approaches the true mean value of *R*_*M*_. Consequently, an estimation of *MI*_*mother*_ can be derived from the mean of *MI*_*front*_*obs*_, ⟨*MI*_*front*_*obs*_⟩, of the child contingency tables by employing the formulas (33) and (35).

From (35), *MI*_*mother*_ is estimated as

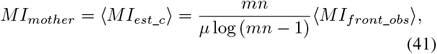

where ⟨*MI*_*est_c*_⟩ is the mean of *MI*_*est*_*c*_ represented as

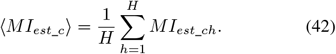

Here, *H* is the number of child contingency tables, while *MI*_*est*_*ch*_ is *MI*_*est*_*c*_ of the *h*-th child contingency table.

By (33), the 95% confidence interval of *MI*_*mother*_ is expressed as

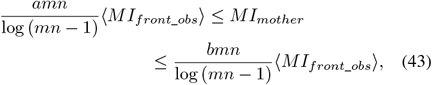

which implies

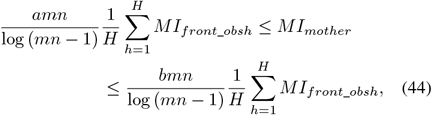

where *MI*_*front_obsh*_ is *MI*_*front*_*obs*_ of the *h*-th child contingency table.

Figure 10 illustrates that the standard error of ⟨*MI*_*est_c*_⟩ is reduced as *m, n* and *H* increase. Concurrently, the estimation error of *MI*_*mother*_ also declines as *m, n* and *H* increase. Consequently, our method is also applicable to meta-analysis, thereby enhancing the precision of estimating the total MI based on the *MI*_*front*_ of the single cell.

## IX. Conclusion and discussion

In this study, we addressed an important issue inherent in the analysis of high-dimensional data, such as genetic data [5, 14, 15, 18]. We developed a novel metric, micromutual information (*MI*_*kl*_), and demonstrated its convergence to total mutual information (*MI*) through a rigorous theoretical framework. This provides an effective method for improving the accuracy of genetic interaction analysis.

We proved Theorem 1, which states that the sum of *MI*_*kl*_ converges to *MI*, as the number of columns and rows tends to infinity. Theorem 1 was extended to Theorems 2 and 3 for multidimensional random variables by using the linear index. We also proved Theorem 4, which asserts that *MI*_*kl*_ asymptotically follows the exponential distribution. Then we defined *MI*_*front*_ as the largest *MI*_*kl*_ and demonstrated that it follows the extreme value distribution. The theorems were supported by the numerical simulations.

Our results demonstrate the utility of the *ab initio* GO method in analyzing complex genetic interactions, particularly in high-dimensional datasets, and show its advantages over traditional methods. We observed that as the number of variable states and dimensionality increase, the estimation of *MI* by a single *MI*_*front*_ becomes more accurate. This aligns with the understanding that genetic interactions occur under specific and constrained conditions, and our approach effectively captures this phenomenon. These findings suggest potential improvements in genetic research methods and offer insights into complex biological systems.

Furthermore, the application of meta-analysis improves the robustness of our estimates, and the integration of multiple datasets allows for more precise *MI* estimation. This complementary approach helps to mitigate the inherent error in estimating *MI* from a single *MI*_*front*_, thereby increasing the utility of our method in complex genetic studies. In contrast to conventional meta-analysis approaches that do not evaluate *P*-values, our method considers the *P*-value of the MI, thereby obtaining *MI* with greater statistical significance.

It should be noted, however, that our work is not without limitations. It remains to be seen whether our approach can be applied to non-genetic, less structured datasets. Further research is needed to adapt the method for broader applications in biostatistics and computational biology. Future work will focus on further developing micromutual information with other data types and extending our theoretical model to a broader range of biological scenarios.

In conclusion, the *ab initio* GO method based on *MI*_*kl*_ and *MI*_*front*_ represents an important step forward in the effort to understand and quantify genetic interactions. It provides a promising new tool for researchers engaged in the rapidly evolving field of genomic sciences.

## Acknowledgment

We are grateful to D. D. Ikeda for assisting numerical simulations.

## REFERENCES

1 Cover, T. M. and Thomas, J. A. (1991). Elements of Information Theory. Wiley, New York.

2 David, H. A. and Nagaraja, H. N. (2003). Order Statistics. Wiley, New York, 3rd ed.

3 Jaynes, E. T. (1957). Information theory and statistical mechanics. Phys. Rev. 106 620–630.

4 Jenkinson, A. F. (1955). The frequency distribution of the annual maximum (or minimum) values of meteorological elements. Q. J. Roy. Meteor. Soc. 81, 158–171.

5 Jeuken, G. S. and Käll, L. (2024). Pathway analysis through mutual information. Bioinformatics 40 btad776.

6 Kinney, J. B. and Atwal, G. S. (2014). Equitability, mutual information, and the maximal information coefficient. Proc. Natl. Acad. Sci. USA 104 501–506.

7 Kraskov, A., Strögbauer, H. and Grassberger, P. (2004). Estimating mutual information. Phys. Rev. E 69 066138.

8 Moon, Y., Rajagopalan, B. and Lall, U. (1995). Estimation of mutual information using kernel density estimators. Phys. Rev. E 52 2318–2321.

9 Mori, T., Kawamura, T., Ikeda, D. D., Goyama, S., Haeno, H., Ikeda, K., Yamaguchi, Y., Shirai, T., Adachi, K., Saito, Y., Horisawa, T., Suzuki, J. and Takenoshita, S. (2023). Influential force: from Higgs to the novel immune checkpoint KYNU. Jxiv 10.51094/jxiv.156.

10 Mori, T. and Kawamura, T. (2023). The equivalence principle of the p-value and mutual information. arXiv http://arxiv.org/abs/2308.14735.

11 Ross, B. C. (2014). Mutual information between discrete and continuous data sets. PLOS ONE 9 e87357.

12 Roulston, M. S. (1999). Estimating the errors on measured entropy and mutual information. Physica D: Nonlinear Phenomena 125 285–294.

13 Shannon, C. E. (1948). A mathematical theory of communications. The Bell System Tech. J. 27 379–424.

14 Song, L., Langfelder, P. and Horvath, S. (2012). Comparison of co-expression measures: mutual information, correlation, and model based indices. BMC Bioinformatics 13 328.

15 The Cancer Genome Atlas Research Network. (2008). Comprehensive genomic characterization defines human glioblastoma genes and core pathways. Nature 455 1061–1068.

16 Witten, E. (2020). A mini-introduction to information theory. La Rivista del Nuovo Cimento 43 187–227.

17 Wyner, A. D. (1978). A definition of conditional mutual information for arbitrary ensembles. Information and Control 38 51–59.

18 Yang, X., Mi, Z., He, Q., Guo, B. and Zheng, Z. (2023). Identification of vital genes for NSCLC integrating mutual information and synergy. Mathematics 11 1460.

